# Towards Zero-Latency Neurofeedback

**DOI:** 10.1101/424846

**Authors:** Nikolai Smetanin, Mikhail A. Lebedev, Alexei Ossadtchi

## Abstract

Neurofeedback (NFB) is a real-time paradigm, where subjects monitor their own brain activity presented to them via one of the sensory modalities: visual, auditory or tactile. NFB has been proposed as an approach to treat neurological conditions and augment brain functions. In many applications, especially in the automatic learning scenario it is important to decrease NFB latency, so that appropriate brain mechanisms can be efficiently engaged. To this end, we propose a novel algorithm that significantly reduces feedback signal presentation in the electroencephalographic (EEG) NFB paradigm. The algorithm is based on the least squares optimization of the finite impulse response (FIR) filter weights and analytic signal reconstruction. In this approach, the trade-off between NFB latency and the accuracy of EEG envelope estimation can be achieved depending on the application needs. Moreover, the algorithm allows to implement predictive NFB by setting latency to negative values while maintaining acceptable envelope estimation accuracy. As such, our algorithm offers significant improvements in cases where subjects need to detect neural events as soon as possible and even in advance.

## 1 Introduction

NFB is a form of biofeedback that allows subjects to perceive and control their own brain activity. Typical NFB settings utilize such recording methods as EEG, magnetoencephalography (MEG) and functional magnetic resonance imaging (fMRI) [1]. Aided by NFB, subjects obtain access to neural signals that they are normally unaware of, and could use these signals in various ways [1], [2], [3], [4]. Steps of NFB operation include extraction of features of interest from neural activity, their transformation into a signal suitable for the user, and delivery of feedback to the subject using one of sensory modalities, for example vision, hearing or tactile sensation. NFB applications emphasize the subjects’ ability to control their own brain activity. Essentially the same paradigm is utilized in brain-computer interfaces (BCIs), but the emphasis of BCI applications is on the ability to communicate with external devices, such as computers and prosthetic limbs [5]. As in NFB systems, sensory feedback is crucial to achieve accurate real-time performance of BCIs.

NFB and BCI paradigms have experienced and explosive development, with many implementations tested in research laboratories and medical clinics. Most importantly, NFB training can result in brain plasticity through a reinforcement learning process [6]. NFB-evoked plasticity can be employed to treat neurological disorders; this is an alternative to pharmacological treatment [7], [8] and cognitive enhancement therapy [9]. Thus, EEG-based BCIs can be used for neurorehabilitation of stroke patients [10], [11]. In this approach, subjects wear an exoskeleton on their hands, and exoskeleton movements are controlled by EEG activity. Alternatively, hand movements can be evoked by functional electrical stimulation of the hand muscles. In both cases volitional cortical activity is synchronized with movement-related proprioceptive feedback, resulting in Hebbian plasticity that eventually repairs the neural circuitry damaged by stroke. Crucially for Hebbian plasticity to occur, there should be a minimal delay between cortical modulations and movements that they trigger.

The efficiency of NFB training depends on many factors that could be classed into spatial, frequency and temporal domains. In the spatial domain, locating the brain region that generates NFB signal allows to target specific functions. To this end, source-space NFB paradigms have been developed, where a spatial filter is derived by either solving the inverse problem [12] or using a spatial decomposition of EEG data [13]. In the frequency domain, specific EEG rhythms are selected to achieve the goals of NFB training. Mathematically, this type of signal is described as a narrow-band stochastic process. This process is usually non-stationary and can be characterized by an envelope that carries information about the instantaneous power of the EEG rhythm of interest.

In the temporal domain, temporal resolution and NFB delay are the key characteristics. The fact that sensory feedback delay affects the efficiency of learning has been recognized in cognitive neuroscience. For example, back in 1948 Grice showed that learning to discriminate complex visual patterns drastically depends on the feedback signal latency [14]. Rahmandad et al. [15] demonstrated that behavioral learning is impaired when feedback delay is unknown. Reduction of frame rate in video playback affects the neuronal mechanisms of feature binding and 3D structure from motion extraction. Increasing feedback delay significantly affects the sense of agency during BCI control [16]. Additionally, a simulation study [17] has demonstrated that feedback delay and temporal blur adversely affect automatic (strategy free) learning.

In a recent study [4] examined the changes in temporal structure of EEG induced by NFB. EEG alpha rhythm was recorded over P4, and mean power of this signal was provided to the subjects via visual feedback. The analysis of the alpha rhythm episodes showed that the subjects were unable to modulate EEG amplitude and episode duration. Instead, they controlled average alpha power by changing the number of alpha episodes per unit time. It has been proposed that treating alpha episodes as discrete events instead of using average power could improve NFB efficiency. In agreement with this idea, [18] showed that the number of beta-band spindles per unit time (and neither amplitude, nor duration) is correlated with cognitive performance in a cued attention task. These findings suggest that onsets and offsets of EEG rhythms are the specifics meaningful events that may be reinforced directly to improve the overal rhythm based NFB.

In the present study, we tackled the issue of NFB delay [1]. To this end, we propose a new method to decrease NFB latency. We first employed an experimental paradigm, where NFB-based learning was affected by the latency of NFB signal. Next, we developed a novel algorithm based on the adaptive optimization of the FIR filter weights. The algorithm estimated the instantaneous power of EEG rhythms at short latency using the dynamic structure present in the EEG signal. This approach can be applied to reducing latency in NFB applications: users could specify the trade-off between the shorter latency and less accurate estimation of the EEG envelope. We also show that even negative latencies are possible, corresponding to predictive NFB.

### 1.1 Feedback latency affects NFB training efficiency

Nine healthy volunteers (5 males and 4 females, aged 22-25) participated in the EEG recordings. One subject was excluded because of insufficient expression of occipital alpha-rhythm power in the eyes-closed as compared to the eyes-open condition. All subjects were right-handed. The experimental procedures were designed and carried out in accordance with the Declaration of Helsinki and approved by National Research University Higher School of Economics Ethical Committee. Written informed consent was obtained from every subject prior to participation in the study. The subjects were paid 300 rubles per hour.

EEG recordings were performed using 32 AgCl electrodes (MCS, Zelenograd) placed according to a modified 10-20-system with the ground electrode at AFz position and reference electrodes on both ears. The impedance for all electrodes was kept below 10 KOhm. The signal was sampled at 500 Hz using the NVX-136 amplifier (MCS, Zelenograd, Russia) and bandpass-filtered in 0.5-70 Hz band. Data acquisition was conducted using NeoRec system and transmitted online via LSL (Lab Streaming Layer). LSL data recording and stimulus presentation was controlled by the in-house NFBLab software. The subjects were seated in a comfortable armchair facing a computer screen. To ensure the presence of a detectable alpha-rhythm, a baseline recording session was conducted prior to the NFB experiment.

We employed P4-alpha NFB. The NFB delay incorporated a 200-ms systematic delay introduced by the signalprocessing pipeline and the hardware. To test different delay values, we introduced additional systematic delay. As outlined in Figure 1, our experiment consisted of feedback *FB_n_* sessions with *n* ∈ [0, 500,1000, *mock*]. The index denotes the artificial delay added to the systematic delay. From the baseline session that preceded NFB experiments, we created individualized spatial filters for on-line rejecting ocular artifacts based on the ICA analysis of the baseline data. The main part of the experiment consisted of four 14-min feedback sessions with 5-min rest intervals. Each NFB session was split into five 2-min blocks separated by 15 s rest intervals. Each NFB session started with a 2-min baseline segment used to estimate the statistics of the alpha-band power dynamics (to be used for standardization purposes in calculating the feedback signal). At the end of each NFB session, we have also collected a 2-min baseline.

**Figure 1:**
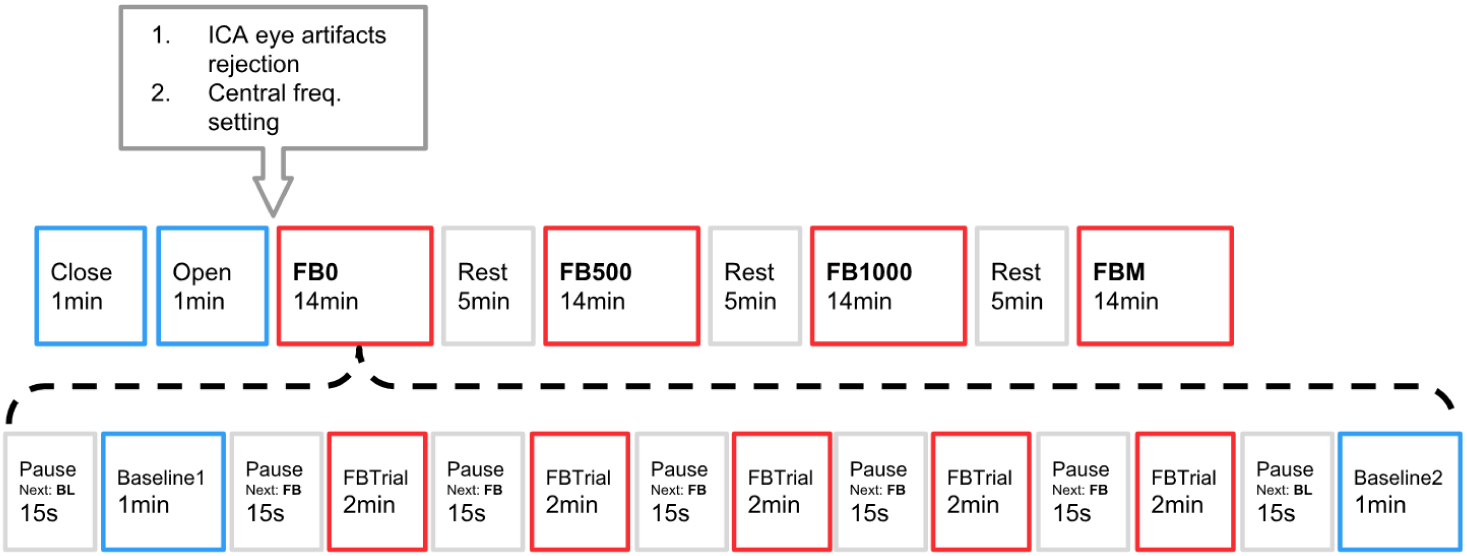
Experiment Design

The sequence of artificial delay values used for each subject was chosen randomly without replacement from a Latin square based design with 4! = 24 blocks. In this non-standard within-subject design we strove to account for the between-subject differences, and therefore each subjected participated in all four types of NFB (including mock-feedback). To counteract the ordering effect we used Latin squares design, so that for each participant the order in which feedback types appeared was different. To further mitigate the ordering effect we introduced 5-min delays between different feedback types. While by the time of writing this report we have collected data from 8 subjects, we plan to increase the number of subjects to 24 to minimize the ordering effects.

The NFB signal depended on the P4-alpha envelope and was visually presented in the form of a “noisy” circle with broken outline on the dark-gray background. The goal of the subject was to smooth the circle outline.

To analyze the effects of the feedback intervention and compare the feedback conditions (14 min sessions) we computed the ratio of the mean 1% quartile of envelope values during each feedback condition to that averaged over all feedback conditions. This quantity was averaged over 8 subjects to yielded the graph in Figure 2.

**Figure 2:**
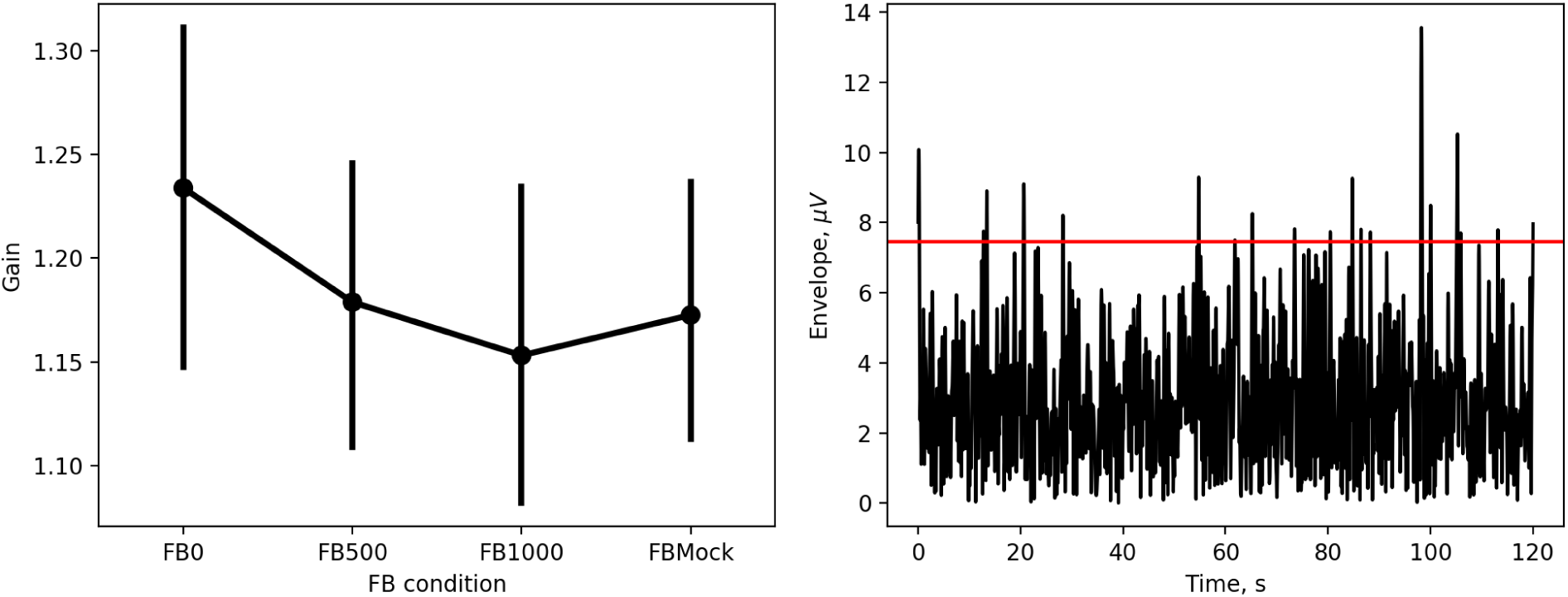
Left panel: alpha power gain vs. feedback latency computed as the average value of the upper 1% of envelope values. Right panel: A representative position of the 99% threshold used to compute the statistics on the Left panel.

We observed that NFB had a greater effect when presented at a shorter latency. With longer delays, NFB effect decreased, and for the longest tested delay, 1200 ms, the results were indistinguishable from those observed during the mock-feedback condition. Although the subjects were unaware of study design, all of them in a subsequent questionnaire reported the sense of agency during the FB_0_ condition and its absence in the other experimental conditions.

Although these results are preliminary, they support the statements regarding the role of NFB delay made in the Introduction and lay out the experimental approach to testing different delay-reducing algorithms.

## 2 Envelope estimation methods

### 2.1 Basic methods

The simplest way to estimate the envelope is to perform low-pass filtering of the rectified narrow-band filtered signal. Formally, this can be represented by the diagram in Figure 3.A. This simple and ubiquitously used approach, however, introduces undesirable delays and leads to the situation where reinforcement for entering into the target brain state (e.g. occipital alpha burst) may be given at the moment when this state has been already abandoned. Attempts to reduce the delay using lower order filters (including their minimum phase versions) result in a rapid deterioration of performance as shown in Figure 5 by the dashed black curve.

**Figure 3:**
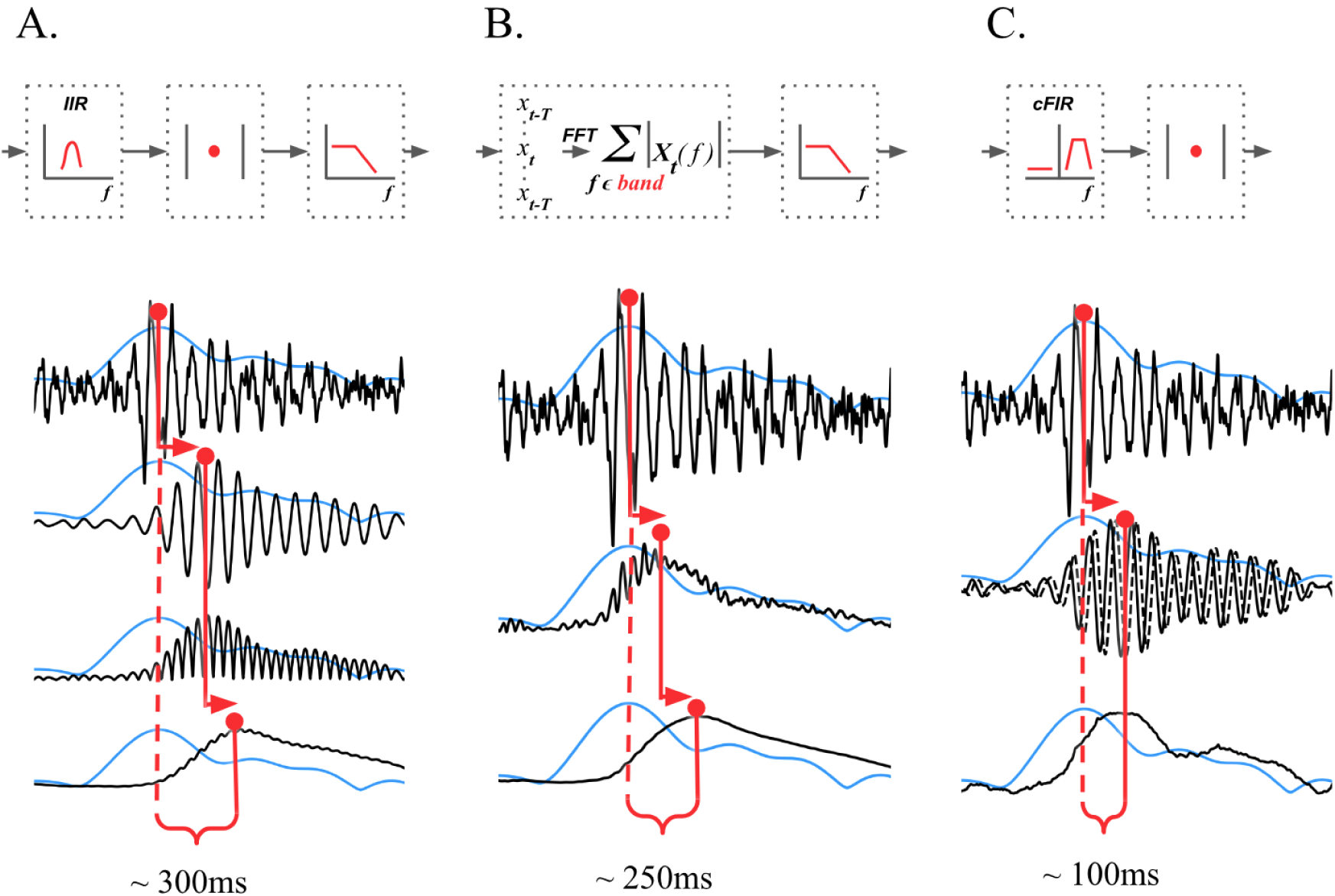
Methods for narrow-band signal envelope estimation: (A) low-pass filtering of the rectified narrow-band filtered signal, (B) STFT with subsequent temporal smoothing of the absolute values of the STFT coefficients, (C) narrow-band analytical signal envelope reconstruction

The other way to obtain the estimate instantaneous band power fluctuations is illustrated in Figure 3.B This method is based on the use of STFT with subsequent temporal smoothing of the absolute values of the STFT coefficients. In addition to the vanilla STFT approach, in order to reduce the latency of the envelope estimate we suggest using mirror reflection of the data segment against the current time moment followed by windowing and FFT. This allows to slightly shorten the delay as compared to the rectifier based technique an yet retain sufficient accuracy of the estimate. Typical delay encountered with this method appears to be around 250 ms. As it is the case with the first method, attempts to reduce the delay also lead to a significant drop in accuracy as illustrated by the solid black curve in Figure 5.

In contrast to these two techniques almost exclusively used in the majority of EEG based NFB paradigm implementations, the family of methods described next delivers significantly better latency-accuracy trade-off. The general idea behind these techniques is based on the use of FIR approximation of Hilbert transform; it is schematically presented in Figure 3.C

### 2.2 Achieving accuracy-latency trade-off in estimation of EEG rhythm power

Fundamentally, the accuracy of envelope estimation is inversely proportional to the length of the interval used to compute an estimate. In contrast to the methods described earlier, the family of techniques presented in this section allows to explicitly specify desired delay value and achieve the best reconstruction accuracy given the specified delay.

The narrow-band brain rhythm signal *s*[*n*] can be represented as real part of analytic signal:

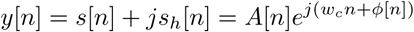

where *s_h_*[*n*] is the imaginary part of the analytic signal, often called “second quadrature” of the original signal *s*[*n*], *w_c_* - central band frequency, *A*[*n*] - envelope, reflecting instantaneous power of the narrow band process, *ϕ*[*n*] - instantaneous phase. Computing the norm of the analytic signal can then be done instantaneously to yield envelope *A*[*n*] of the original signal *s*[*n*].

To build analytical signal *y*[*n*] corresponding to the original narrow-band signal *s*[*n*] which is a part of noisy broad-band signal *x*[*n*] one can apply a narrow-band Hilbert transform filter. Frequency response of the combined filter can be defined as:

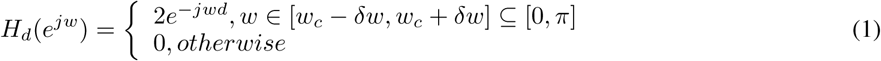

where *δω* is half of passband width, *d* - group delay in samples. For any finite *d* this filter is non-causal and cannot be applied in real time. To reconstruct analytical signal causally one can find complex-valued finite impulse response filter (cFIR) which approximates the ideal frequency response *H_d_*(*e^jw^*) [19]. The desired filter b can be found by solving least square optimization problem defined in different ways.

The first and most straightforward approach (denoted F-cFIR) that follows least squares filter design strategy is to find the complex valued vector of cFIR filter weights b that minimizes the *L*_2_ distance between cFIR filter frequency response and the ideal response *H_d_* in the frequency domain:

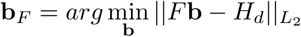

where *F* is the DFT transformation operator. This simple approach does not take into account the structure of EEG signal and can be further improved.

Following optimal filter design ideas we can use power spectral density *X* of the input signal *x*[*n*] as weights. We thus formulate the weighted frequency domain least squares design technique (denoted WF-cFIR) via the following optimization problem:

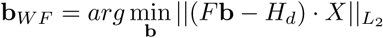

This method allows to exploit the regularities and hidden relationships between rhythmic components in EEG signal.

Figure 4 illustrates these two approaches. The delay *d* corresponds to the slope of the phase response within the passband and can be set to an arbitrary value. Then, the optimization procedure aims at finding such complex vector of FIR filter coefficients b that both phase response and amplitude response are approximated sufficiently accurately. We do so in the frequency domain by analytically solving least squares or weighted least squares problems. Conceptually, the described approach allows to accommodate different weights to the errors in approximating amplitude and phase responses of the resultant filter. In this case, however, we would have to perform an iterative optimization in order to find the optimal FIR filter weights vector b.

**Figure 4:**
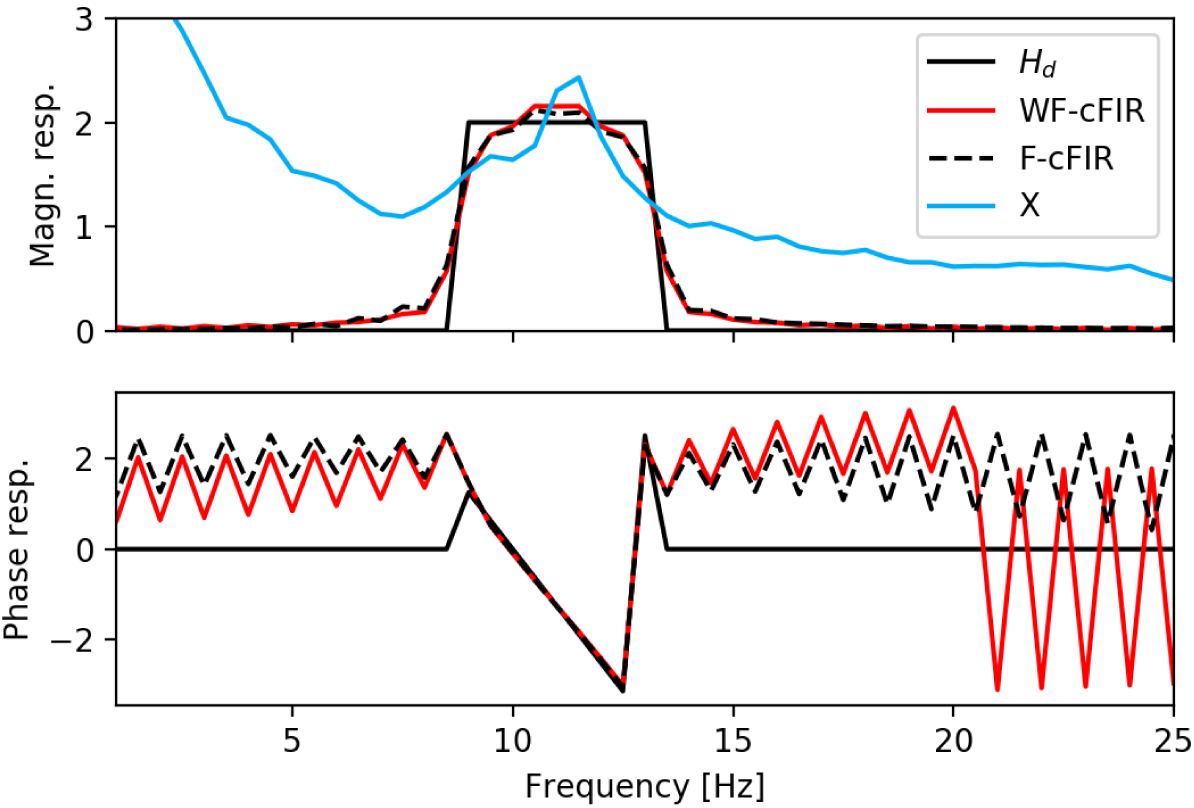
Ideal analytical signal extraction filter amplitude and phase frequency response (solid black line) and its WF-cFIR (red line) and F-cFIR (dashed black line) approximation. Blue line corresponds to power spectral density of the input signal

The last approach from this family (denoted T-cFIR) is based on minimization of the squared distance in the time domain between the complex delayed ground truth signal *y*[*n* – *d*] and the filtered signal *x*[*n*] * b[*n*]:

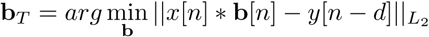

where * is the convolution operator. In this case during the training stage the ground truth signal *y_d_* is obtained non-causally from the training dataset via ideal zero-phase Hilbert transformer (1). According to Parseval’s theorem this approach is equivalent to the WF-cFIR approach, however in contrast to the frequency domain formulation it allows in a straightforward manner to add amplitude dependent weights in order to achieve better accuracy in approximating the bursts of activity.

### 2.3 Results

We compared the described methods using the data recorded in the NFB sessions conducted in 8 subjects. As the ground truth, we used non-causally computed envelope as the absolute value of the analytic signal obtained via frequency domain Hilbert transform. We then calculated the correlation coefficient for the envelope obtained causally by each of the five methods with the ground-truth sequence, and plotted it as a function of the corresponding delay. For the T-CFIR and WF-CFIR approaches the statistics was computed using cross-validation procedure, based on the contiguous testing segments. To vary delay for the first two techniques that do not allow to explicitly specify it we adjusted filter order which led to changes in the group-delay time.

The results of this comparison are summarized in Figure 5. As we can see, the cFIR based family of methods results in significantly better performance and yields high correlation between the true and estimated envelopes even for zero-latency. Most intriguing is the fact that with these methods we can introduce negative latencies and perform prediction of the envelope values. Also note, as shown in the top-right panel of 5, this approach applied to the white noise signal delivers significantly weaker performance than in the situation with the real EEG signal.

**Figure 5:**
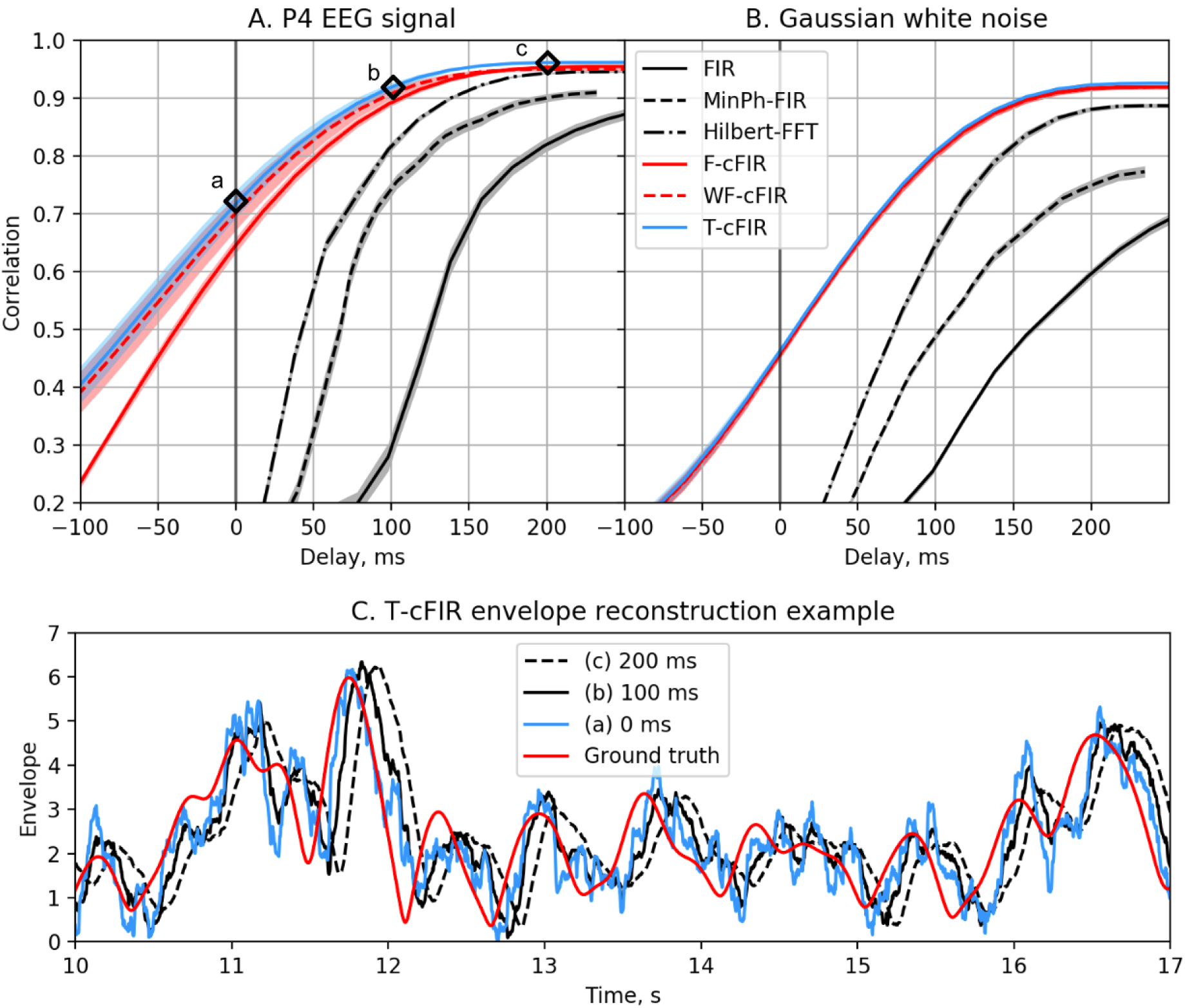
Correlation coefficient for the envelope obtained causally by each of classic (black curves) and novel (colored curves) filters with the ground-truth sequence vs. filters group delay for real EEG P4-channel data (A) and white noise signal (B). Example of T-cFIR envelope reconstruction for 0, 100 and 200ms delays (C)

Next, we conducted a numerical experiment in which we used the data from a subject involved in a discrete NFB paradigm where the subject received NFB based on the envelope computed with rectification of the narrow-band filtered signal followed by low-pass filtering. This method incurs a 300 ms delay and results in the feedback signal arriving toward the end of the downslope of the alpha-burst, see Figure 6. We then experimented with the WF-cFIR approach for different delay parameter *d* and plotted the real-time alpha-envelope obtained by averaging time-locked to the feedback arrival moment. This way, we could appreciate the true delay including the hardware and lower level software details that amounted to approximately 100 ms. Note that even a 100-ms additional processing delay led to the NFB signal arriving close to the downslope of the alpha-burst for *d* = 0. The proposed method is capable of predictive behavior, as evident from the results for *d* = –50*ms*. In this case NFB is delivered around the upward slope of the alpha-burst.

**Figure 6:**
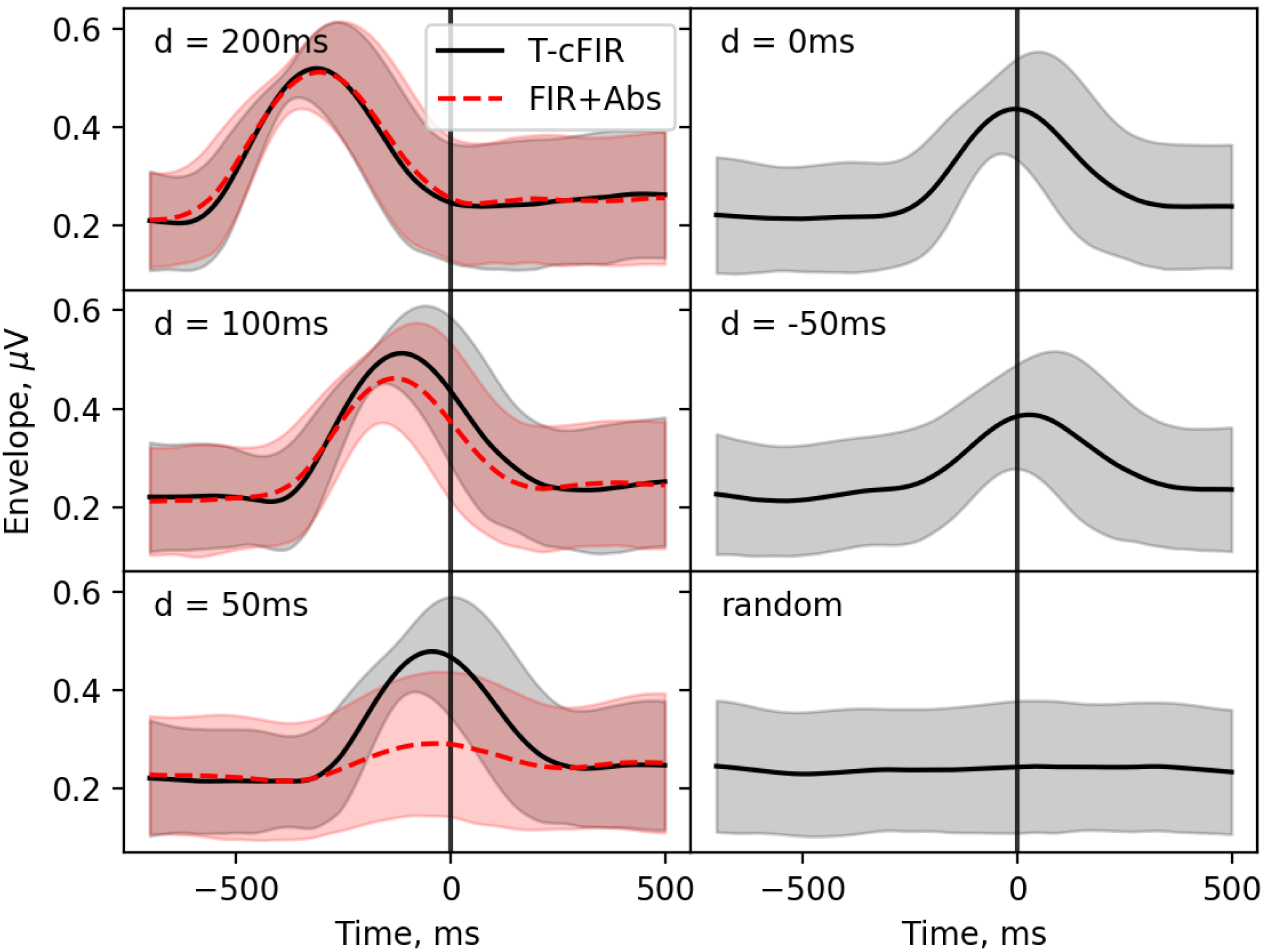
Envelope of the P4-channel averaged by feedback onset moments as registered by the screen mounted photo-sensor for different delay values. Note, that even 100 ms delay leads to the feedback signal arrival at the downslope of the alpha-burst. At the same time, the proposed method proves to be capable of predictive behavior and for *d* = –50*ms* allows to deliver the feedback on the up-slope of the alpha-burst taking into the low level hardware delays.

## 3 Discussion

We have shown that the standard techniques for estimating instantaneous power of EEG rhythms, such as methods based on rectification of narrow-band filtered signal and the algorithms employed in the STFT, incur significant delays, which may potentially hinder the efficiency of NFB. With these methods, reinforcement for correct performance arrives more than 300 ms past the transient neural event meant to be reinforced. Such a delay is especially clear when changes in narrow-band signals are considered as discrete events. For discrete estimates of EEG states, NFB generated with the standard techniques often arrives after the state transition is over. Such imperfect timing of neural activity and associated NFB may interfere with NFB tasks in unwanted ways.

To address this problem we suggested a series of methods based on the least-squares filter design. The methods allow to directly control the accuracy versus latency trade-off for the assessment of narrow-band signals. Our results showed that this approach significantly reduced the latency resulting from EEG signal processing. Thus, spectral density weighted cFIR technique allowed to generate zero-latency NFB that tracked EEG-rhythm power with sufficient accuracy. As evident from comparison of the plots in the left and right panels of Figure 5, short-latency processing was possible because of the presence of specific EEG patterns that allowed signal extrapolation into the future. (When applied to Gaussian random process signal these methods performed significantly worse.)

The approaches presented here can be further extended by employing more sophisticated decision rules capable of extracting the hidden structure from the data. Convolutional neural networks [20] being a natural extension of the approach presented here, hold a significant promise to further improve the accuracy of zero-lag envelope estimation.

The improved delay values came at a cost of less accurate envelope estimation. For example, an attempt to achieve zero or even negative delay may result in significant errors in the envelope estimation, leading to NFB malfunction. Therefore, the optimal trade-off between the envelope estimation and NFB latency is the question that needs to be addressed for each particular application. An example of envelope estimate obtained for *d* = 0 condition overlayed onto the true envelope is shown in Figure 5.C.

Yet, we suggest that the efforts to build a zero- or even negative-latency NFB systems will eventually pay off because they lead to an implementation of predictive NFB paradigm. To achieve the true predictive scenario, improvements need to be made not only to the signal processing algorithms used for feature extraction but also to the hardware employed for signal acquisition, as well as low-level software that handles EEG data transfer from the acquisition device to the computer memory buffer. It is worth considering specialized systems based on FPGA programmable devices to eliminate the uncertain processing delays present in computer operating systems that are not designed to operate in real-time.

Additional consideration should also be given to the physiological aspects of the sensory modality (or modalities) used to present the NFB. For instance, it is known that the visual pathways, although very informative [16], are significantly slower compared to tactile feedback. Thus, combining different sensory modalities could improve temporal characteristics of NFB while keeping information transfer rate high.

## Acknowledgments

This work was supported by the Center for Bioelectric Interfaces of the Institute for Cognitive Neuroscience of the National Research University Higher School of Economics, RF Government grant, ag. No. 14.641.31.0003.

## References

[1] Sitaram R., Ros T., Stoeckel L., Haller S., Scharnowski F., Lewis-Peacock J., Weiskopf N., Blefari M., Rana M., Oblak E., Birbaumer N., and Sulzer J. Closed-loop brain training: the science of neurofeedback. Nature Reviews Neuroscience, 2016.

[2] Sterman M., MacDonald L., and Stone R. K. Operant control of the eeg alpha rhythm and some of its reported effects on consciousness. Altered states of consciousness, 1969.

[3] Sterman M., MacDonald L., and Stone R. K. Biofeedback training of the sensorimotor electroencephalogram rhythm in man: effects on epilepsy. Epilepsia, 1974.

[4] Ossadtchi A., Shamaeva T., Okorokova E., Moiseeva V., and Lebedev M. Neurofeedback learning modifies the incidence rate of alpha spindles, but not their duration and amplitude. Scientific Reports, 2017.

[5] Wolpaw J. et al. Brain–computer interfaces: principles and practice. Oxford Univ. Press, 2012.

[6] Lawrence E. J. et al. Self-regulation of the anterior insula: Reinforcement learning using real-time fmri neurofeedback. Neuroimage, 2014.

[7] Lofthouse N., Hendren R., Hurt E., Arnold L. E., and E. Butter. A review of complementary and alternative treatments for autism spectrum disorders. Autism research and treatment, 2012.

[8] Coben R., Linden M., and Myers T.E. Neurofeedback for autistic spectrum disorder: a review of the literature. Applied Psychophysiology and Biofeedback, 2010.

[9] Zoefel B., Huster R. J., and Herrmann C. S. Neurofeedback training of the upper alpha frequency band in eeg improves cognitive performance. Neuroimage, 2011.

[10] Frolov A., Kozlovskaya I., Biryukova E., and Bobrov P. Robotic devices in poststroke rehabilitation. Zhurnal Vysshei Nervnoi Deyatelnosti Imeni I.P. Pavlova, 2017.

[11] Ang K. K., Chua K. S., Phua K. S., Wang C., Chin Z. Y., Kuah C. W., Low W., and Guan C. A. Randomized controlled trial of eeg-based motor imagery brain-computer in-terface robotic rehabilitation for stroke. Clinical EEG and neuroscience, 2015.

[12] Congedo M., Lubar J.F., and Joffe D. Low-resolution electromagnetic tomography neurofeedback. IEEE Transactions on Neural Systems and Rehabilitation Engineering, 2004.

[13] White D.J., Congedo M., and Ciorciari J. Source-based neurofeedback methods using eeg recordings: training altered brain activity in a functional brain source derived from blind source separation. Front. Behav. Neurosci, 2014.

[14] Grice G. R. The relation of secondary reinforcement to delayed reward in visual discrimination learning. Journal of experimental psychology, 1948.

[15] Rahmandad H., Repenning N., and Sterman J. Effects of feedback delay on learning. System Dynamics Review, 2009.

[16] Evans N., Gale S., Schurger A., and Blanke O. Visual feedback dominates the sense of agency for brain-machine actions. PLoS ONE, 2015.

[17] Sulzer JS Oblak E.F., Lewis-Peacock JA. Self-regulation strategy, feedback timing and hemodynamic properties modulate learning in a simulated fmri neurofeedback environment. PLoS Comput Biol, 2017.

[18] Shin H., Law R., Tsutsui S., Moore C., and Jones S. The rate of transient beta frequency events predicts behavior across tasks and species. eLife, 2017.

[19] Wei Rong Lee, Lou Caccetta, Kok Lay Teo, and Volker Rehbock. Optimal design of complex fir filters with arbitrary magnitude and group delay responses. IEEE Transactions on Signal Processing, 2006.

[20] Schirrmeister R.T., Springenberg J.T., Fiederer L., Glasstetter M., Eggensperger K., Tangermann M., Hutter F., Burgard W., and Ball T. Seldeep learning with convolutional neural networks for eeg decoding and visualization. Hum Brain Mapp, 2017.

